# Cortical rhythms are modulated by respiration

**DOI:** 10.1101/049007

**Authors:** Detlef H. Heck, Samuel S. McAfee, Yu Liu, Abbas Babajani-Feremi, Roozbeh Rezaie, Walter J. Freeman, James W. Wheless, Andrew C. Papanicolaou, Miklós Ruszinkó, Robert Kozma

**Affiliations:** Dept. of Anatomy & Neurobiology, University of Tennessee Health Science Center, Memphis, TN, 38163; Dept. of Pediatrics, University of Tennessee Health Science Center and Le Bonheur Children’s Hospital, Memphis, TN, 38105; Department of Molecular & Cell Biology, Division of Neurobiology, University of California at Berkeley, Berkeley CA 94720-3206 USA; Rényi Institute of Mathematics, Hungarian Academy of Sciences, Budapest, Realtanoda u., Hungary; Dept. of Mathematical Sciences, University of Memphis, Memphis, TN 38152

## Abstract

The brain generates oscillatory neuronal activity at a broad range of frequencies and the presence and amplitude of certain oscillations at specific times and in specific brain regions are highly correlated with states of arousal, sleep, and with a wide range of cognitive processes. The neuronal mechanisms underlying the generation of brain rhythms are poorly understood, particularly for low-frequency oscillations. We recently reported that respiration-locked olfactory bulb activity causes delta band (0.5-4 Hz) oscillatory neuronal activity in the whisker sensory (barrel) cortex in mice. Furthermore, gamma oscillations (30 – 100Hz), which are widely implicated in cognitive processing, were power-modulated in synchrony with the respiratory rhythm. These findings link afferent sensory activity caused by respiration directly to cortical rhythms associated with cognitive functions. Here we review the related literature and present new evidence to propose that respiration has a direct influence on oscillatory cortical activity, including gamma oscillations, and on transitions between synchronous and asynchronous cortical network states (marked by phase transitions). Oscillatory cortical activity, as well as phase transitions, has been implicated in cognitive functions, potentially linking respiratory phase to cognitive processing. We further argue that respiratory influence on cortical activity is present in most, and possibly in all areas of the neocortex in mice and humans. We furthermore suggest that respiration had a role in modulating cortical rhythms from early mammalian evolution. Early mammals relied strongly on their olfactory sense and had proportionately large olfactory bulbs. We propose that to this day the respiratory rhythm remains an integral element of dynamic cortical activity in mammals. We argue that breathing modulates all cortical functions, including cognitive and emotional processes, which could elucidate the well-documented but largely unexplained effects of respiratory exercises on mood and cognitive function.

## Introduction

The rhythm of respiration is one of the fundamental rhythms of life. It is inextricably linked to olfaction, a sensory modality of great importance to the evolutionarily earliest mammals, which possessed correspondingly large olfactory bulbs (Rowe et al., 2011). Neurons in the olfactory bulb, at the first central processing stage of odors, rhythmically increase and decrease their firing rates with each breath, even in the absences of odors (Adrian, 1950; Phillips et al., 2012). Thus, via olfactory bulb output, respiration generates a continuous rhythmic neuronal input to the piriform (olfactory) cortex and drives respiration-locked firing in piriform cortical neurons (Fontanini and Bower, 2005; Fontanini et al., 2003). Respiration-locked oscillations have also been observed in areas that receive direct projections from the piriform cortex, such as the prefrontal cortex and the hippocampus (Nguyen Chi et al., 2016; Tsanov et al., 2014; Yanovsky et al., 2014). Here we review published evidence, as well as present new data supporting the view that respiration influences three distinct aspects of cortical neuronal activity - possibly in all cortical lobes. We suggest that the functional consequence of respiration-locked cortical activity is a rhythmic modulation of sensory perception, and of motor and cognitive functions.

Published results, as well as new (unpublished) data from our laboratory show that respiration does not only affect neuronal activity at the respiratory frequency but also causes phase-locked modulation of the power of gamma band oscillations (30 – 100 Hz) (Ito et al., 2014) and of the timing of phase transitions (Freeman, 2015; Kozma and Freeman, 2016). Based on these findings we suggest that respiration is a source of variability of cortical activity and consequently, cortical function. The modulation of the power of gamma oscillatory activity has been widely implicated in the performance of a broad range of sensory and cognitive tasks (Canolty et al., 2006; Herrmann et al., 2010; Kay et al., 2009) and the occurrence of phase transitions in cortical activity has been associated with the occurrence of sudden insight or understanding (“aha” moments) (Freeman, 2004b, 2015; Kozma and Freeman, 2008; Kozma and Freeman, 2016).

Taken together, the evidence we summarize here may require a drastic revision of our current conceptions of the physiology of respiration, which are based on gas exchange and olfactory sensation. We suggest a radically new view of respiration by proposing that respiration directly modulates cognitive brain function by synchronizing neuronal activity across large areas of neocortex. Since cognitive processes in turn affect respiratory behavior, respiratory modulation of cognitive function indicates the presence of an intimate, life-long interaction between mind and body. As a consequence, the intentional modulation of respiration may, if fully understood, become a powerful tool for the manipulation of cognitive function. At the same time, abnormal respiratory rhythms may reflect and/or cause cognitive impairment. This in turn suggests that long-term observation of respiratory behavior, similar to ambulatory EKG, may be a powerful physiological and psychological diagnostic tool.

## Respiration locked oscillations in mouse and human cortex

We recorded local field potentials (LFPs) and spike activity in various areas of the mouse cortex while simultaneously monitoring respiratory behavior. LFP and spike activity was then analyzed to determine whether activity was modulated in phase with respiration. LFP activity was averaged aligned with the end of the expiration cycle. Whether LFP activity was significantly modulated by respiration was determined using bootstrap statistics (Diaconis and Efron, 1983; Efron and Tibshirani, 1993). Surrogate LFP averages for bootstrap statistics were created by randomly shuffling respiration times and recalculating LFP averages aligned on shuffled respiration times. This process was repeated >100 times to determine *p* = 0.05 significance boundaries (see methods section for details).

Spike activity was analyzed by cross correlating spike times with end-of-expiration times. Again, bootstrap statistics were used to determine whether spike rate was significantly modulated in phase with respiration. Figure 1 shows respiration locked averages of LFP activity and spike respiration cross-correlations for 5 different recordings sites, which include the prefrontal, somatosensory, primary motor and visual cortex, and the olfactory bulb. Respiration locked modulation of LFP and spike activity in the sensory whisker barrel cortex (asterisk in Fig. 1) has been shown previously (Ito et al., 2014).

**Figure 1:**
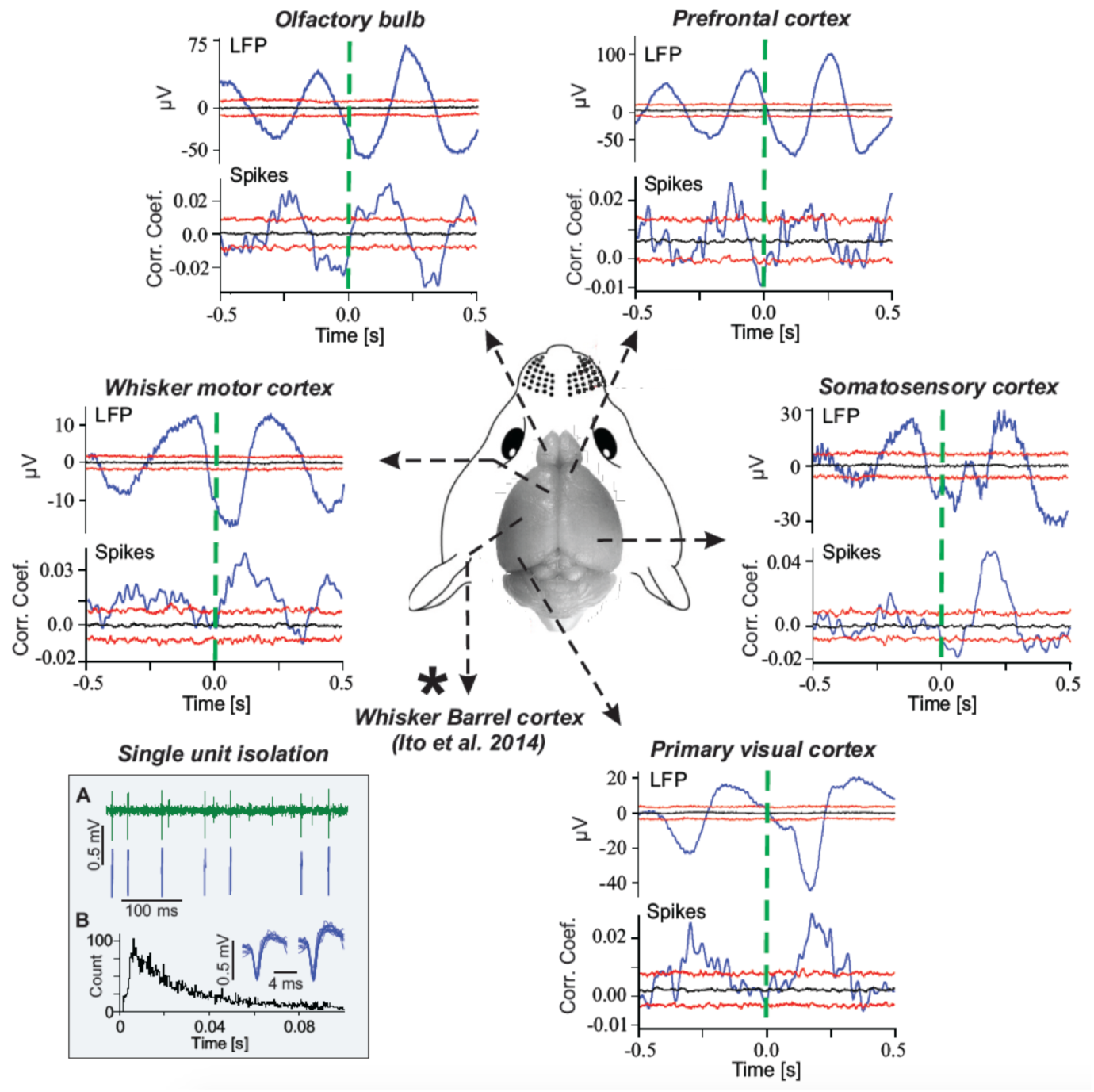
Spike and LFP activity in different areas of the neocortex of awake mouse are phase locked to respiration. Blue lines in histograms are cross-correlations. Dashed green vertical lines mark the end of expiration. Horizontal black and red lines represent the predicted median (black) and the 5 and 95%ile boundaries (red) of the surrogate data obtained by random shuffling (Bootstrap statistics). Square insert at the bottom left shows an example single unit recording in primary motor cortex. A in insert) Raw spike recording with sorted single unit spikes shown below. B in insert) Inter-spike interval (ISI) histogram showing the typical shape of a singe unit with effects of refractory period. Traces above ISI show overlays of the first and last 10 spikes recorded over 300 seconds.

LFP and spike activity at each recording site were significantly modulated in phase with the rhythm of respiration indicating that in mice, respiration is a major contributor to cortical delta oscillations. In a previous study we have shown that delta oscillations in the mouse sensory whisker barrel cortex are driven mostly by respiration-locked olfactory bulb sensory activity (Ito et al., 2014). The spike activity of olfactory bulb output neurons (mitral and tufted cells) is unfailingly modulated in phase with respiration, even in the absence of odors (Adrian, 1950; Phillips et al., 2012). This is likely due to the fact that olfactory sensory neurons not only detect chemical substances or odors, but are also mechano-sensitive and respond the pressure changes associated with the movements of air through the nasal cavity during inspiration and expiration (Grosmaitre et al., 2007). Removing the olfactory bulb eliminated respiration locked activity in the whisker barrel cortex almost entirely (Ito et al., 2014).

It is very likely, however, that breathing-related sensory inputs from other sources also contribute to respiration-locked cortical oscillations. Amongst those are chemical sensors in the blood vessels detecting blood CO2 levels, stretch receptors in the lungs, mechano-sensitive cells in the upper airways and stretch receptors of the diaphragm and intercostal muscles (Belvisi, 2003; Davenport and Vovk, 2009; Finger et al., 2003; Schelegle, 2003; Widdicombe, 2009). Most of this interoceptive sensory activity is represented in the anterior insular cortex (Craig, 2002, 2009). In addition to the sensory activity caused by respiration, there are also projections from the brain stem to the thalamus (Carstens et al., 1990; Krout et al., 2002). Some of these projections likely provide respiration-locked input to the thalamus (Chen et al., 1992) introducing a non-sensory respiration-locked input to the thalamo-cortical circuit. How exactly different pathways contribute to cortical respiration-locked activity remains to be determined. At the current state of knowledge it seems that in highly olfaction-reliant rodents the olfactory bulb is a main driving force. In humans, however, judging by the small size of the olfactory bulb compared to the rest of the brain, olfaction is likely to play a lesser role in driving respiration-locked cortical activity.

Recordings of cortical activity in humans were obtained from epilepsy surgery patients, implanted with subdural grid electrodes for the purpose of electrocorticographic (ECoG) monitoring and localization of epileptiform activity. The signals recorded with subdural grid electrodes correspond to LFP recordings obtained in mice and were analyzed and interpreted in the same way. Respiration-triggered averaging of ECoG signals from 3 patients showed a significant modulation of ECoG in phase with the respiratory cycle at most recording sites. Sites showing respiration locked neuronal activity were found in all lobes covered by the subdural grid, namely the temporal, the parietal and the frontal. Examples of ECoG averages showing respiration-locked modulation over the three lobes are shown in Fig. 2. Most electrode locations were in areas of the human brain not involved in olfactory processing. At the time of the writing of this manuscript, recordings from the occipital cortex were not available. Recording sites for subdural grid electrodes are determined by medical considerations and none of the patients in this study required occipital placement of electrodes. We calculated respiration-triggered averages of raw ECoG signals for all recordings sites and found significant respiration-locked modulation of cortical activity at more than half of the recording sites. Together, the results obtained in humans and mice show that respiration, likely due to respiration locked sensory inputs, synchronizes neuronal activity across large areas of neocortex, at the rhythm of respiration.

**Figure 2:**
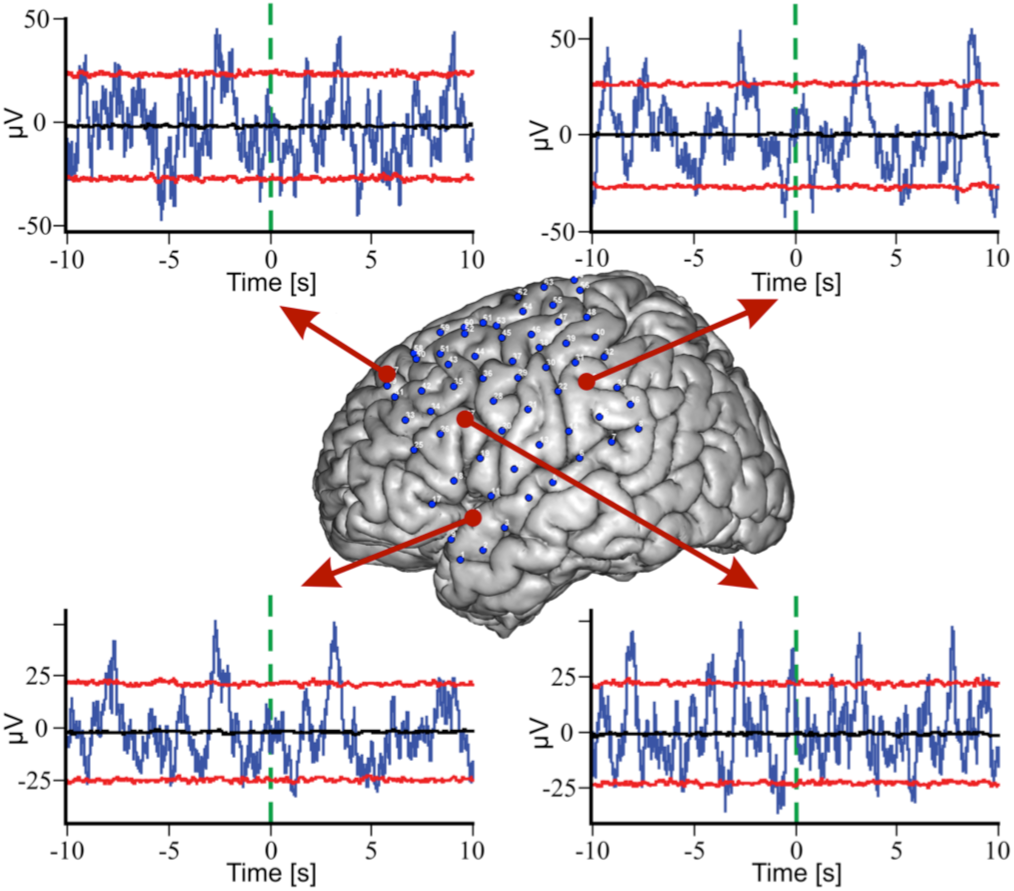
Electrocorticographic (ECoG) activity in different lobes of human neocortex is phase locked to respiration. Four graphs showing averages of ECoG signals recorded at the cortical locations indicated by the red arrows. Averages were aligned on the times of expiration onset (green dashed vertical line). Blue lines in graphs represent ECoG averages; black and red lines represent the median, 95%ile and 5%ile boundaries of the surrogate data distribution, respectively (see Methods). The image of the brain in the center has blue dots marking ECoG recording sites. Arrows link recording sites in the frontal, parietal and temporal lobes with the plots of expiration-triggered ECoG averages calculated for that site. In all three lobes ECoG activity was significantly modulated in phase with breathing (p<0.05), resulting in peak ECoG activity at ~3 sec before and after expiration onset. The inter-peak interval corresponds to the patient’s average respiratory interval of about 6 sec.

## Respiration locked modulation of gamma power

The cortical rhythms generated at different frequencies are often interdependent. A common observation is that high-frequency oscillations show rhythmic increases and decreases in amplitude, which are phase locked to the rhythm of a lower frequency oscillation. A well-described example of such phase-amplitude coupling is the modulation of gamma oscillation (30-100 Hz) amplitude in phase with the slow theta (4-8 Hz) oscillations (Canolty et al., 2006). The strength of phase locking between gamma power and theta phase varies with the specific task performed. Learning and working memory in rodents are accompanied by changes in theta-gamma phase-amplitude coupling (PAC) in the prefrontal cortex and hippocampus (Li et al., 2012; Tort et al., 2009), which suggests that theta-gamma PAC reflects cognitive processes. We have shown that respiration modulates gamma oscillatory power in the mouse whisker barrel cortex (Ito et al., 2014). Here we asked if this finding extends to other cortical areas and other frequency bands. To address this question we performed a PAC analysis based on analytic amplitude taken from a Hilbert transform of the band-passed filtered LFP signals, to determine which, if any, frequencies of cortical oscillations showed respiration-locked amplitude modulation. In mice we found significant modulation of gamma amplitude at all cortical sites tested.

**Figure 3:**
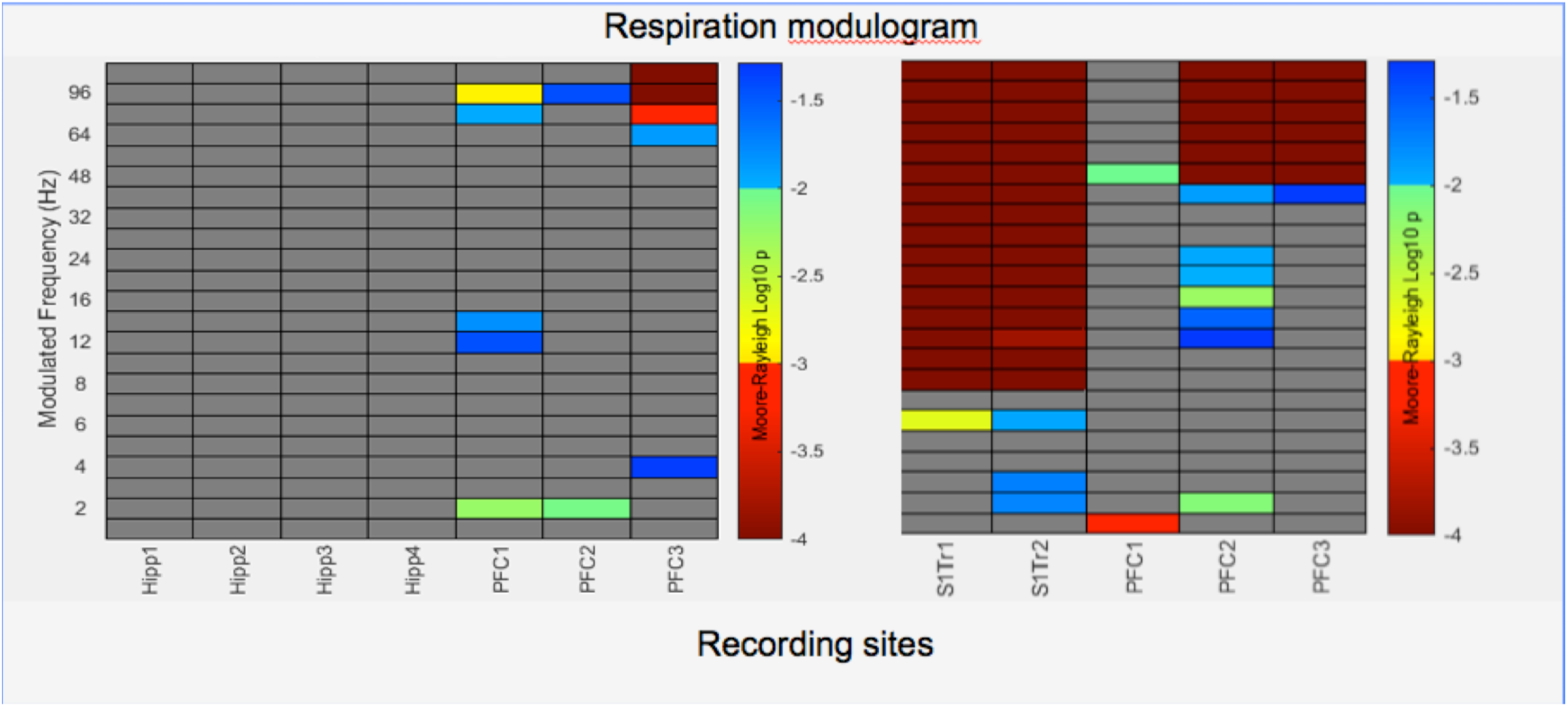
Phase-amplitude coupling of respiration with neuronal (LFP) oscillations in mouse hippocampus, prefrontal cortex and somatosensory cortex. Panels show modulograms for two different mice. ***Left panel:*** LFP recordings were performed at four sites in the hippocampus (Hipp1-4) and at three sites within the prefrontal cortex (PFC1-3). Significant respiration-locked power modulation (pseudo colors code Moore-Rayleigh *p* value) were seen only in the prefrontal cortex and were most significant for gamma band oscillations. ***Right panel:*** LFP recordings were performed at two sites in the trunk area of the somatosensory cortex (S1Tr1-2) and at three sites in the prefrontal cortex (PFC1-3). Phase amplitude coupling occurred across a broad band of frequencies (8 – 100 Hz) at both sites in the somatosensory cortex. In the prefrontal cortex phase amplitude coupling was most strongly expressed in the gamma range and only observed at two of three recordings sites. One recording site also showed respiration-modulated power in the theta/beta frequency range.

The same analysis of human ECoG data revealed widespread synchronous modulation of gamma amplitude across all three cortical lobes covered with grid electrodes in three patients. In one patient (MN, Fig. 5), almost all recording sites showed significant modulation in a narrow gamma frequency band. In another patient, gamma power modulation was limited to the inferior frontal lobe (MS, Fig. 5). In the third patient (RV, Fig. 5) respiration-locked gamma modulation was scattered and seen at only few recordings sites. At this point we do not understand what determines whether a cortical site shows respiration-locked modulation of gamma activity. But we asked whether this modulation had a preferred respiratory phase during which the probability for gamma-power increase was highest. We used Moore-Rayleigh statistics to estimate the phase during which gamma power had the highest probability of increased amplitude. The probability of increased gamma power was highest during the resting phase of respiration, i.e. the period between two breaths. This finding was consistent across both patients with consistent gamma activity modulation (patients MS and MN). Mice do not have as pronounced a resting phase between breaths as humans do but consistent with the findings in humans the preferred phase for increased probability of high gamma power in mice was around the time of transition from expiration to inspiration.

**Fig 5.**
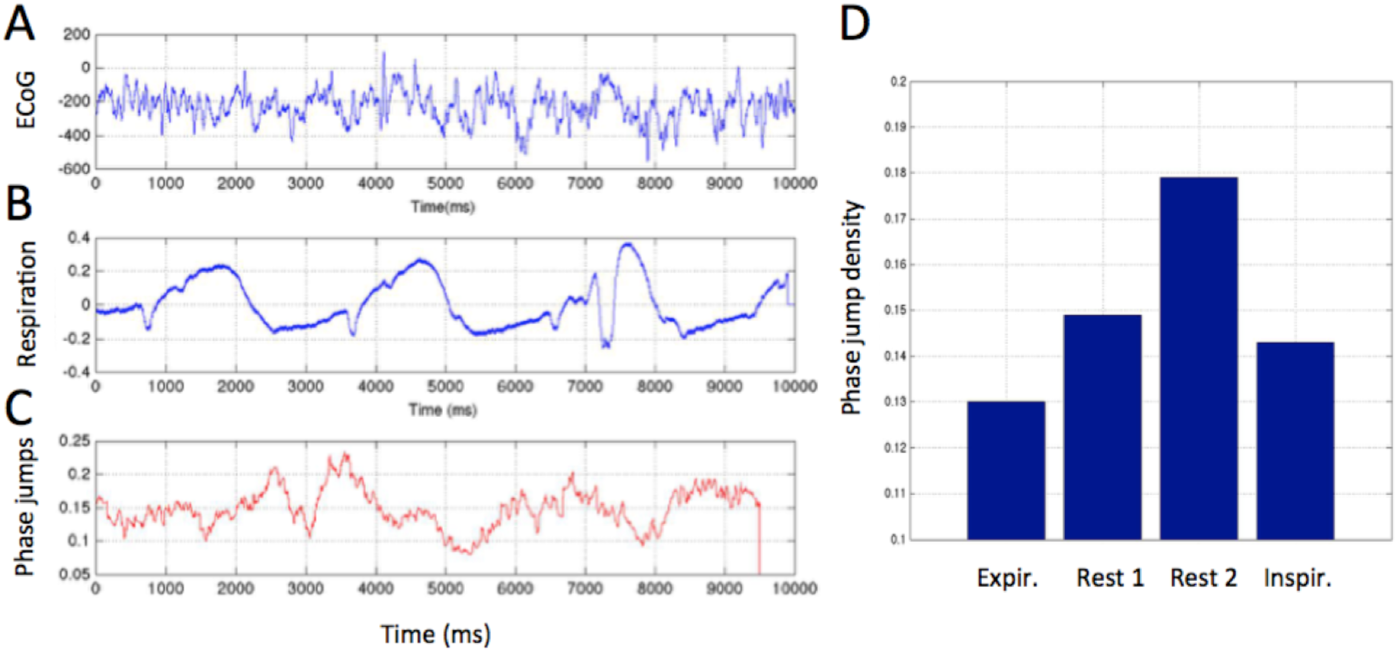
Illustration of the Hilbert analysis method using 58 traces of raw ECoG data and the simultaneously recorded respiratory signal. ***A-C)** Three subplots showing **A)** a single raw ECoG signal, **B)** raw respiratory signal, and **C)** the integral effect of phase jumps across 58* **subdural grid** *electrodes. Phase jumps were determined as the discontinuities of the phase of the analytic signal exceeding a certain threshold over the gamma band* (Freeman, 2009). *A threshold of 60 degrees was used here. An average value of phase jump 0.1 means that there are about 6 phase jumps at a given time, while phase jump value 0.2 means twice as many phase jumps (around 12). The individual raw ECoG signal in A) does not exhibit an obvious correlation with the respiratory signal (B). However, the integral phase discontinuity value (C) is phase-locked to the respiratory signal. **D)** Phase jump value distribution across four discrete phases of the respiratory cycle divided into expiration, first and second half of the resting phase (Rest 1, Rest 2), and Inspiration. Phase jump activity is significantly increased during the resting period*.

**Figure 4:**
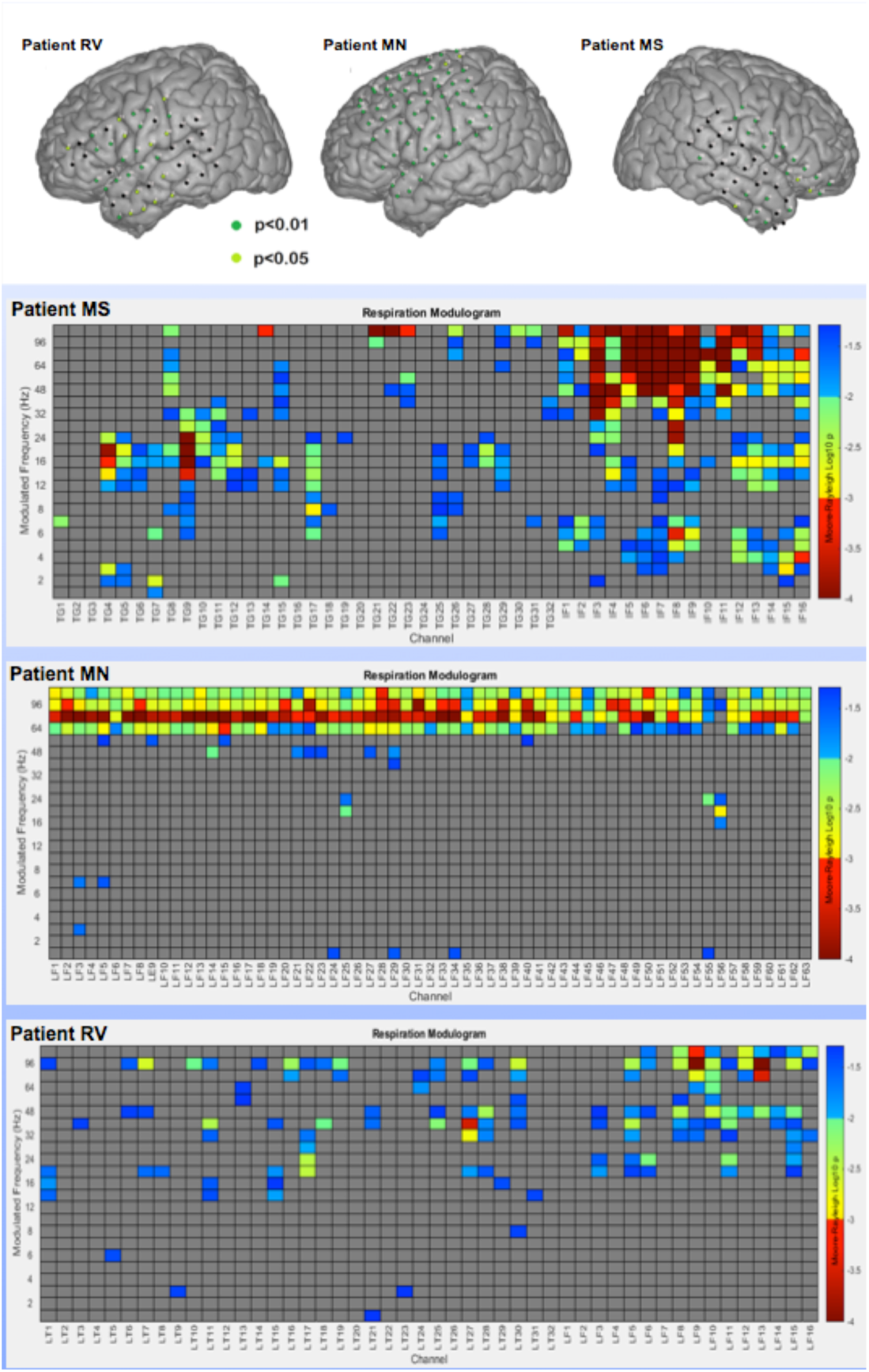
Phase-amplitude coupling of respiration with neuronal oscillations in the human neocortex. ***Top panel:*** Subdural electrode locations, shown on a rendering of the standard Montreal Neurological Institute (MNI) brain atlas, in 3 patients with epilepsy who underwent a Phase II epilepsy surgery evaluation. ***Bottom panels:*** modulograms for each patient showing frequencies (y-axis) with significant respiration-locked power modulation (pseudo colors code Moore-Rayleigh *p* value) for each ECoG electrode (x-axis).

## Respiration locked timing of phase jumps

In the case of rapidly changing, non-stationary signals, Hilbert analysis can provide useful insight into neuronal processes. Hilbert analysis is based on determining the analytic signal and its instantaneous frequency, which can be used to describe phase synchronization effects in neuronal populations. Sudden changes in phase synchronization across cortical areas have been linked to cognitive processing (Kozma and Puljic, 2015). In the Hilbert approach, one determines the analytic amplitude and analytic phase of complex-valued Hilbert-transformed signals at various frequency bands (Kozma et al., 2012). The analytic phase is predominantly a continuous function of time, which may, however, show sudden discontinuities (phase jumps). These phase jumps mark brief, transient periods, which have been suggested to have cognitive significance, possibly associated with moments of sudden insight (“aha” moments) (Freeman, 2004a).

Here we use Hilbert analysis to identify relationships between respiration and phase discontinuities within the ECoG signals. Figure 5 illustrates the results of such analysis using a raw LFP signal from an array of 58 surgically implanted, subdural ECoG electrodes from an epilepsy patient. Figure 5a shows an example of the raw ECoG/LFP signal, while Fig. 5b is the respiratory signal, with inspiration and expiration represented by increasing and decreasing voltage, respectively. The respiratory signal has a triphasic sequence consisting of inspiration, expiration and rest, with a total duration of around 3 seconds in the example shown. The average number of large phase jumps (exceeding 60 degrees) in the analytic phase across the array is plotted in Fig. 5C. There is statistically significant correlation between the number phase jumps and the respiratory phase. This is emphasized in Fig. 5D, where the average phase jump values are depicted for the expiration, rest (part 1&2), and inspiration phases.

## A simple graph theory model of respiratory modulation of cortical activity

#### INSERT BOX: BASIC GRAPHY THEORY TERMINOLOGY

For general concepts of graph theory, see, e.g., (Erdos and Renyi, 1960; Watts and Strogatz, 1998; Newman and Watts, 1999; Benjamini and Berger, 2001; Albert and Barabasi, 2002; Bollobas et al., 2007). Graph theoretical approaches have been used successfully since the early 2000’s to model properties of brain networks (Sporns et al., 2005; Reijneveld et al., 2007; Bollbas et al., 2009; (Gallos et al., 2012; Reijneveld et al., 2007; Turova and Villa, 2007). A review of the literature on graph theory approaches to brain structure and dynamics is given, e.g., in (Kozma and Freeman, 2016).

Here we summarize the graph terminology used in this study for describing the model for the respiratory modulation of cortical activity.

- Graph: A mathematical object G(n, M), with a collection of n vertices (nodes) with M edges (connections) between some pairs of the vertices.
- Path: A path between two nodes is the sequence of edges starting in one node and ending in the other. The number of nodes along the path measures path length.
- Distance: Distance between two nodes is the length of the shortest path between them.
- Diameter: Diameter of a graph G is the length of the longest distance between any pairs of its edges.
- Connectedness - A graph is connected if it contains a path between any two vertices.
- Degree of a node - The number of edges connected to that node.
- Random graph - Graphs when the vertices and or edges are selected in some random way. In the Erdos-Renyi random graph (Erdős and Rényi, 1960), the numbers n of vertices and M of edges are given, and the random graph is chosen uniformly from among all graphs with vertex set {1,2,…n} and M edges.
- Giant component - A component of the graph, which contains a constant (high) fraction of the nodes.
- Geometric graph - Has its nodes placed in the (Euclidean) space. Here we consider 2-dimensional planar geometry.

An example of a geometric graph defined on the 2-dimensional plane is shown in Fig. 6. Note that the vertices of this graph are placed on a square lattice. Each vertex has a short edge (undirected) to its direct neighbor, all together 4 edges. In addition to these short edges, the diagram shows some additional long edges that are selected randomly between two vertices that are not direct neighbors on the square lattice grid.

**Figure 6.**
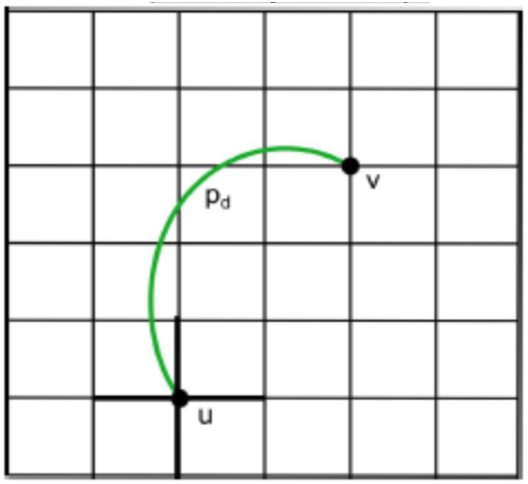
Illustration of a geometric graph defined over the 2-dimensional square lattice of size 7x7. *The 4 local edges of node u are marked with bold black lines. There is a long edge between node u and v shown in green. The probability of the long edge is given by p_d_, as defined in the model description of the main text*.

END OF BOX: BASICS GRAPH THEORY FOR BRAINS

In order to determine biologically plausible conditions that would allow a neocortex-like network to support experimentally observed properties of large-scale oscillations, we explored the parameter space of a simple graph theoretical model of the neocortex. Here we focus on the respiratory modulation of gamma power, while other observations, such as the timing of phase jumps being phase-locked to respiration, will be the objective of future studies. Our results show that the influence of respiratory phase on the modulations of the gamma power can be reproduced in our graph theoretical model within biologically meaningful parameters describing network connectivity and excitation-inhibition balance.

When using a graph theoretical approach, we view brains as large-scale networks, with some nodes and edges connecting the nodes (see box “Basics of graph theory for brains”). According to the adopted population model, the nodes are not representative of individual neurons, but of populations of thousands of neurons (up to about 10,000) which correspond to granules of neurons (Freeman, 1975; Kozma and Freeman, 2009). Edges between the nodes indicate connections between neuronal populations. Depending on the goal of the study, the connections may correspond to structural or functional links between neuronal populations. In our case, connections mean that the populations influence each other. Most of the edges are relatively short reflecting the fact that neuronal connections via axons are concentrated around their dendritic arbor. There are however, longer connections extending beyond the direct neighborhood of the nodes. The number of such longer connections is relatively low, nevertheless, they play crucial role in the propagation of neural activity between distant cortical areas (Kozma and Puljic, 2015).

We consider graphs in a two-dimensional planar geometry, corresponding to the cortical sheet (Freeman et al., 2009). In mathematical terms, we define a graph using a 2-dimensional square lattice of *N* × *N* points folded into a torus, and denote it as 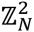. The points of the lattice give the vertices of the random graph. The edges of the generated graph 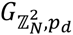 include the regular edges of the lattice, and some additional edges selected randomly. The random edges are selected between two nodes with probabilities *p*_*d*_ that are defined with respect to the lattice graph distance *d* of vertices to be joined. The distribution of the long edges reflects the biological fact there are many more short connections than long ones in the neural tissue:

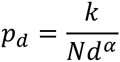

where α is the exponent of a power-law long edge distribution; for simplicity, here we use the value *α* = 1. The regular edges of the lattice model the local connections between neural nodes, while the additional long edges model non-local connections mediated by long axons.

On the constructed graph, we consider the propagation of activity in the presence of excitatory and inhibitory effects. As the starting step, a full mathematical analysis of the activation process on 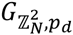 in the presence of purely excitatory nodes has been given. The main result in (Janson et al., 2016) determines all stable and unstable fixed points, which provide the critical initial probability values *p*_*k*_ indicating phase transitions. That is, in case of initializing the process below critical values, the system will die out; all nodes eventually will become inactive. On the other hand, in case of initializing the process above critical values, the system will completely recover. Next we consider models with inhibition. The results (Kozma et al., 2016) show the presence of parameter regions with convergent behavior (fixed point), as well as multiple fixed points (stable and unstable), indicating oscillations. The oscillations inherent in coupled excitatory-inhibitory (E-I) populations describe the generation of gamma oscillations in the cortical neuropil.

We modeled respiratory effects by periodic perturbations in the form of varying probability of the initial activation of the populations. In order to investigate the effect of respiratory modulation of gamma power in mouse brains, we modeled an intrinsic oscillation in the E-I population, which was 8-10 times faster than the respiratory rhythm. This corresponds to the fact that respiration at theta rates in the mouse (~3-4 Hz) is about 10 times slower than the dominant gamma rate of cognitive processing.

Our simple model, which reproduced respiration-induced modulation of gamma fluctuations, has several key parameters: the density of long connections (*λ*), the proportion of excitatory nodes (*ω*), and the magnitude of the driving respiratory modulation (RA). *λ* = 0 means the absence of long connections; ω changes between 0 (pure inhibition) and 1 (pure excitation). In the introduced mean-field model, one has to solve the coupled system of fixed-point equations for the density of excitatory (*χ*) and inhibitory (*y*) populations

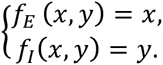

*ƒ*_*E*_ in the first equation describes the transition of the densities of active excitatory nodes, while *ƒ*_*I*_ in the second equation models the transition of densities of inhibitory nodes. The actual forms of these functions are quite complex consisting of several components, for details, see (Janson et al., 2016). For illustration, we show a relatively simple component in mean-field approximation, describing the probability that an active inhibitory node remains active following an update step:

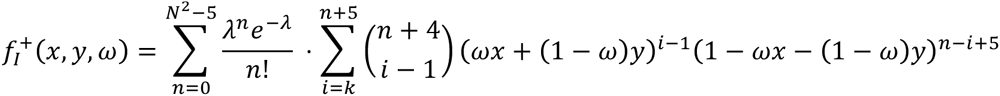

Here *k* is a threshold parameter of the model describing the likelihood that the activity of the neighbors will impact the activity of a given node. In other words, *k* is the sensitivity of the nodes to input effects. The above equation incorporates the mathematical result (Janson et al., 2016; Kozma et al., 2015) that the degree distribution of the nodes in graph 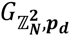 can be approximated well with Poisson (Barbour et al., 1992) statistics for large system size (*N*).

Numerical evaluations show the parameter regions where limit cycles are present (Fig. 7, deep blue region). Dampened oscillations are observed in the light blue regions, while pale color shows rapid convergence to fixed-point regime. By properly tuning the parameters of the model, we can produce sustained oscillations describing the gamma regime of excitatory-inhibitory populations.

**Figure 7.**
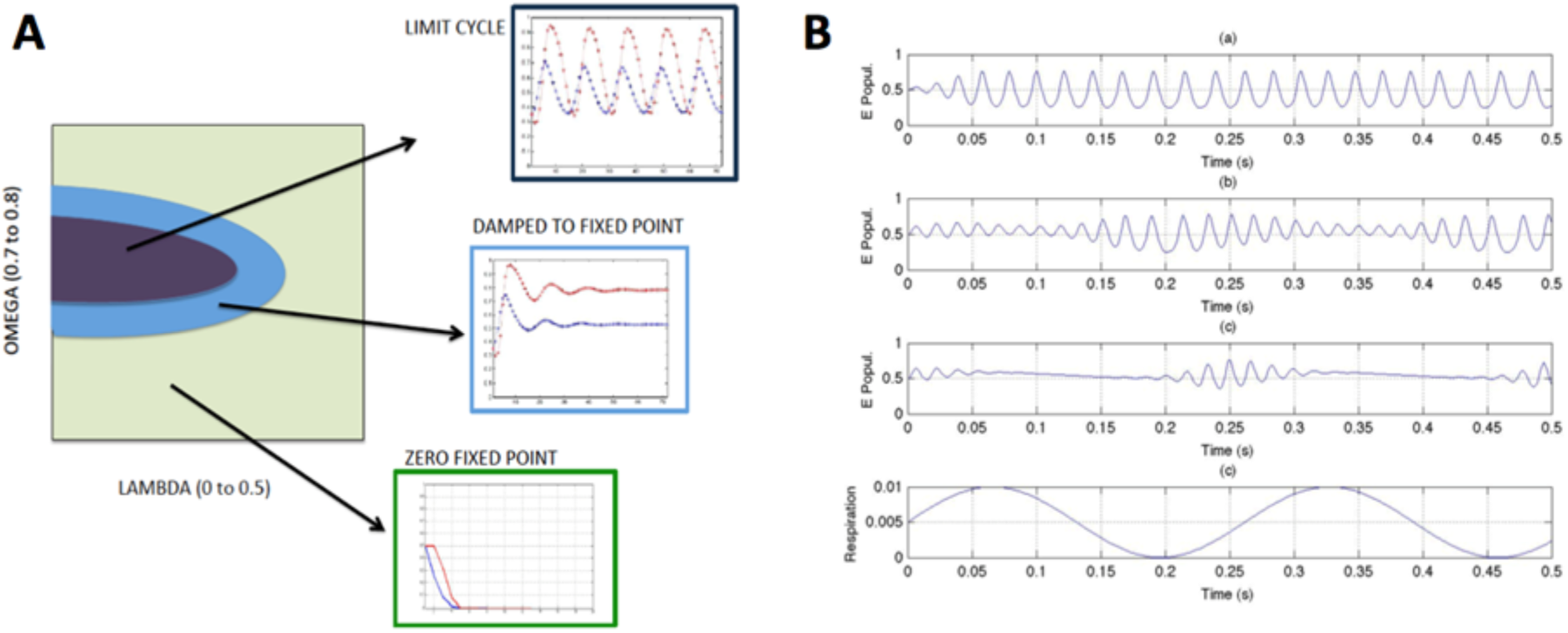
Results of calculations using graph theory models of coupled excitatory-inhibitory populations. *(a) Phase diagram with parameter regions of limit cycle, nonzero fixed point, and zero-fixed point regimes; (b) Illustration of the phase-locked amplitude modulation of the gamma oscillations (of excitatory population) in response to input (respiratory) perturbations of increasing amplitude (RA); w=0.75, lambda=0.01; (a) RA=0.001; (b) RA=0.02; (c) RA=0.03; (d) shape of the respiratory sinusoid signal*.

Our results show increased gamma activity during the segment of the respiratory cycle with increasing amplitudes (inhalation). On the other hand, gamma activity is significantly reduced when the respiratory signal is in the decreasing stage (exhalation). The above results are in agreement with observations made during the ECoG experiments with mice. We conclude that our graph-theoretical model using sine modulation of soft gamma oscillators is able to reproduce and interpret basic properties of the experimentally observed phase-locking between respiration and gamma modulation.

## Discussion

Ongoing fluctuations of neuronal activity have been widely considered random noise that introduces variability into neuronal processing, which is averaged out by the brain through integration of population activity (Georgopoulos et al., 1986; Lee et al., 1988; Maynard et al., 1999; Shadlen and Newsome, 1994). It is now understood that the fluctuating component of cortical activity consists of highly structured activity patterns, including oscillations at various frequencies, that modulate evoked neuronal responses (Arieli et al., 1996; He, 2013; Poulet and Petersen, 2008) and affect sensory perception (Boly et al., 2007; Linkenkaer-Hansen et al., 2004; Palva et al., 2013; Sadaghiani et al., 2009; Vinnik et al., 2012). Ongoing cortical activity is attributed to proprioceptive and interoceptive inputs as well as to intrinsically generated activity, which could be related to mental processes (Deco et al., 2011; Fox and Raichle, 2007).

Here we argue that respiration is a long overlooked, major contributor to neuronal activity that causes rhythmic modulations of at least three major aspects of cortical activity. Based on published and new evidence from both mice and humans, we argue that respiration causes 1) slow oscillations of cortical single unit and LFP activity that follow the species-specific respiratory rhythm, 2) modulation of the power of gamma oscillations in phase with respiration, and 3) the timing of phase transitions to preferentially occur during a specific phase of the respiratory cycle.

### Physiological mechanisms of respiration-locked oscillations

Respiration creates both conscious and unconscious streams of rhythmic sensory inputs to the brain. Consciously accessible sensations of normal, unobstructed breathing include odor perception, the mechanical and thermal sensation of air flowing through nose, mouth and upper airways, and the proprioception of movements of the chest and abdomen. Unconscious sensory signals caused by respiration include interoceptive signals from the lungs, diaphragm and internal organs, which represent the mechanical consequences of respiratory movements, and the chemosensitive signals from the cardiovascular system, which represent breath-by-breath fluctuations of CO2 and Oxygen levels in the blood. The sensations and brain activity patterns associated with hunger-for-air (Liotti et al., 2001; Macey et al., 2005) are not considered here, as they represent an emergency response not related to normal, unobstructed breathing.

There are also indirect ways cortical areas receive respiration-locked sensory input. Eye movements, for example, have been shown to be temporarily phase-locked to respiration during sleep (Rittweger and Popel, 1998) as well as in the awake state (Rassler and Raabe, 2003). Recently, Ito and colleagues reported saccade related changes in LFP oscillation power in four frequency bands, including gamma, in primates freely viewing their environment (Ito et al., 2013). This suggests that the retinal flow associated with eye movements causes a modulation of power in visual cortical oscillations that is partially correlated with respiration. The auditory cortex receives auditory input related to respiration caused by the sound of air flowing through the nose or mouth. Finally, neurons in the brain stem project broadly to thalamic nuclei (Carstens et al., 1990; Krout et al., 2002). These projections likely provide respiration-locked input to the thalamus (Chen et al., 1992), introducing a non-sensory respiratory rhythm to the thalamo-cortical network.

Despite this multiplicity of cortical afferents that carry respiration-locked neuronal rhythms, the influence of respiration on cortical activity can be subtle. Typically, it cannot be assessed without simultaneously measuring respiration and brain activity and then relating the neuronal activity to respiration. Such simultaneous measurements are rarely performed, which is one likely reason why the influence of respiration on cortical activity has remained undetected. A notable exception is a recent study of the effects of sleep disordered breathing (SDB) on cortical oscillatory activity(Immanuel et al., 2014). Immanuel and colleagues showed that the average power of the EEG signal decreased during inspiration and increased during expiration, in a frequency band and sleep stage dependent manner, in both healthy subjects and subjects suffering from SDB (Immanuel et al., 2014).

While there are many sources of respiration-locked activity, the olfactory system deserves special attention, because early mammals relied strongly on their olfactory sense and had proportionately large olfactory bulbs (Rowe et al., 2011). Furthermore, neuronal oscillations, particularly gamma oscillations, are a universal element of odor processing in animals as far removed from joint evolutionary ancestors as mammals and insects (Kay, 2015). Even though in primates the olfactory sense lost the prime importance it has for most other mammals in favor of vision (Gilad et al., 2004), EEG studies comparing nasal and oral breathing of room air found that nasal breathing elicited significantly different patterns of EEG activity than mouth breathing (Lorig et al., 1988; Servit et al., 1977). This is in line with our findings related to nasal air flow in mice (Ito et al., 2014) and suggests that olfactory bulb activation by nasal airflow also contributes to human respiration-related EEG activity. A small group of researchers have envisioned the possibility of respiration influencing large-scale brain activity via the olfactory system. Freeman and colleagues performed pioneering studies on the influence of respiration through olfaction on the rabbit brain (Eeckman and Freeman, 1990; Kay and Freeman, 1998). Effects of theta-modulation of saccadic signals have been described as visual sniffing (Kozma and Freeman, 2001). Fontanini and Bower speculated that olfactory bulb respiration-locked oscillations in rodents may propagate through the entire cortex (Fontanini and Bower, 2006). However, none of these earlier studies anticipated that respiration would modulate the power of gamma oscillations and the timing of network state transitions (phase-slips), which suggests a direct link between respiratory phase and cognitive brain processes.

Respiration related sensory activity during unobstructed breathing mainly reaches three areas of the cortex: 1) the olfactory cortex and surrounding areas receive olfactory bulb input, 2) the somatosensory cortex receiving input from mechanoreceptors of chest, the abdominal skin and muscles stretched and moved by respiration and 3) the insular cortex receives input from chemoreceptors and mechanoreceptors in the lungs, diaphragm and internal organs. Our recordings in mice and humans show that respiration-locked activity propagates from these primary sensory areas to parts of the cortex that do not receive direct respiration related sensory inputs, such as primary motor areas and the primary visual cortex. A likely mode of propagation is through the cortico-cortical network itself, possibly involving also cortico-thalamic connections. However, the anatomy of axonal connections within the parabulbar and limbic areas suggest a number of subcortical regions and neuromodulator systems may also be influenced by respiration-driven sensory input. For example, widely projecting serotonergic and cholinergic neurons within the rat basal forebrain have been shown to rhythmically discharge in phase with respiration (Linster and Hasselmo, 2000; Manns et al., 2003; Mason et al., 2007). Stimulation of cholinergic neurons in particular is associated with increased neocortical gamma oscillations (Cape and Jones, 2000), a mechanism that might contribute to the respiration-locked modulation of gamma power described here.

If respiratory sensory activity propagates in this way through the cortex one would predict that sensory activity from other modalities propagates in similar ways, modulating neuronal activity in other cortical areas. Evidence that this is indeed the case is abundant in the literature. For example, Peterson and colleagues used voltage sensitive dye imaging in awake mice to study cortical activity in the primary whisker motor cortex in response to mechanical stimulation of a single mystacial whisker (Ferezou et al., 2007). Whisker stimulation caused the expected topographically localized sensory response in the primary whisker sensory (barrel) cortex. Within about 60 ms, however, the initially localized sensory response had propagated throughout almost the entire cortex, including the contralateral hemisphere (Ferezou et al., 2007).

While performing EEG recordings in humans, McDonald et al. showed that an auditory stimulus that was otherwise irrelevant to the task, elicited a neuronal response in the contralateral primary visual cortex (McDonald et al., 2013). Following up on these findings, Feng and colleagues later showed that the visual-cortical response evoked by auditory stimulation had functional significance. The amplitude of the auditory-evoked visual cortical response predicted the probability of a correct response to a visual discrimination task (Feng et al., 2014). Neill and colleagues showed that the behavioral state of a mouse, resting or running on a treadmill, had dramatic effects on visual cortical activity and visually evoked responses (Niell and Stryker, 2010). We attribute those differences to the fact that a walking or running mouse experiences massing somatosensory inputs to the cortex, which also spread to the visual cortex and modulate neuronal responses to visual stimuli. Taken together these findings show that exteroceptive and interoceptive sensory inputs shape ongoing neuronal activity and sensory processing in the cortex and that sensory activity propagates through the cortical network beyond the boundaries of primary sensory areas.

### Physiology and Modeling of respiratory modulation of higher-order cortical activity

It is possible to link the existence of respiration-locked cortical oscillations directly to respiration-related sensory inputs to the cortex. However, the phase-locked modulations of gamma power and of the timing of phase transitions cannot be directly explained by rhythmic respiration-locked sensory inputs entraining cortical activity. These forms of respiratory influence on higher-order patterns of cortical activity seem to be emergent properties that could depend on a number of factors including the balance of excitation and inhibition and their patterns of connectivity within the cortical network. Experimental results using various brain-imaging tools evidence the presence of synchronization effects. These results lead to the postulation of the existence of functional links between cortical regions, e.g., when the correlation between cortical areas exceeds a threshold (Bonifazi et al., 2009; Honey et al., 2010; Kim et al., 2013; Sporns et al., 2005; Stam et al., 2007).

To investigate the processes leading to transient synchronization effects in the cortex, we used a simple graph theory model inspired by cortical network architecture and functions, with a biologically appropriate balance of excitatory and inhibitory neurons and mix of short-and long-range connections. Expanding on previous work (Gallos et al., 2012; Reijneveld et al., 2007; Turova and Villa, 2007), the present study reveals important properties between biologically observed effects of respiration on gamma power modulation reproduced by the graph theory model, suggesting that the cortical network itself is sufficient to modulate gamma power in phase with the respiratory rhythm. This is not to say that other factors, such as cortico-thalamic interactions or the action of neuromodulators have no role but future research will have to determine the nature of their involvement.

### Functional implications

Each of these forms of cortical activity appear to have different functions. Oscillatory rhythms that are phase-locked to respiration may help to synchronize large portions of the cortical network and create a temporal alignment for slower processes. The calming effect of controlled, slow and deep breathing could be due to this respiration-locked synchronization of activity across large areas of cortex, an EEG activity pattern commonly observed during meditative states (Dillbeck and Bronson, 1981; Gaylord et al., 1989). ADDITIONAL EVIDENCE OF RESPIRATION-LOCKED SYNCHRONIZATION OF CORTICAL OSCILLATORY ACTIVITY COMES FROM A STUDY OF EEG ACTIVITY DURING MEDITATION WITH FORCED ALTERNATE NOSTRIL BREATHING, WHICH CAUSED AN INCREASE IN INTERHEMISPHERIC BETA COHERENCE (STANCAK AND KUNA, 1994).

Oscillations of neocortical neuronal activity in the gamma (30 – 100 Hz) frequency range, have been strongly implicated in affective and cognitive brain functions such as attention (Fries et al., 2001; Laufs et al., 2003; Tallon-Baudry, 2004), sensory perception (Engel et al., 2001; Gould et al., 2012; Tallon-Baudry, 2003), decision making (Gould et al., 2012; Kay and Beshel, 2010; Nacher et al., 2013; Siegel et al., 2011; van Vugt et al., 2012; Wyart et al., 2012), problem solving (Sheth et al., 2009) and memory formation (Chauvette et al., 2012; Marshall et al., 2006; Tort et al., 2009). Respiratory modulation of gamma oscillation power is thus likely to influence neuronal processes associated with gamma oscillations. There is currently no data to support or refute this hypothesis, as monitoring respiration during cognitive tasks is not routinely done. However, a similar principle seems to be responsible for the dynamic modulation of gamma oscillatory power and synchronization in the primate visual system. Lowet and colleagues recently reported that rhythmic synchronization of gamma oscillations between areas V1 and V2 and increases in gamma power are phase-locked to microsaccades (Lowet et al., 2015). Similar to respiration, rhythmic microsaccades generate a visual sensory input that modulates ongoing activity in visual cortical neurons (Martinez-Conde et al., 2013). Thus, the ability of sensory inputs to synchronize neuronal activity within and between cortical networks and to modulate the power of gamma oscillations is likely not limited to respiration. Inputs from other sensory modalities might have similar effects.

Detailed analysis of rabbit and human intracranial ECoG signals have revealed discontinuities in the analytic phase (which are also called phase transitions) determined by Hilbert analysis (Freeman, 2015; Freeman et al., 2006; Freeman and Rogers, 2002). Experiments with rabbits trained using classical conditioning paradigm showed that these phase sips have cognitive relevance (Freeman, 2004b; Kozma and Freeman, 2008). Namely, after delivering the conditioned stimulus, the occurrence and duration of the qualifying phase transitions correlates with the stimulus, suggesting that phase transitions can be viewed as marker of the cognitive activity (classification) performed by the rabbits.

Phase-slips mark the transitions of cortical network states from synchronous to asynchronous and vice versa. Schölvinck and colleagues observed that variability of neuronal responses in the primary visual cortex to repeated identical stimuli was caused by large scale network activity, which was more variable when the network was in a synchronized state vs. an asynchronous state (Schölvinck et al., 2015). Thus, the fact that phase-slip timing is coupled to respiration suggests that respiration modulates cortical network state-transitions and consequently the variability of sensory processing. Sharp wave ripples in the hippocampus, a high-frequency (100 - 200 Hz) oscillatory event that synchronizes large populations neurons (Csicsvari et al., 1999), may be related to phase transitions. Recently, the role of sharp wave ripples has been outlined as prominent synchronous population patterns in the hippocampus, which affect wide areas of the cortex (Buzsaki, 2015). Sharp wave ripples influence cognitive functions, action planning, and potentially may influence creative thoughts (Ritter et al., 2012). Buzsaki emphasizes the complex interaction between sharp waves ripples and theta rhythm on modulating gamma oscillations, and they may play a role in the observed theta modulation of gamma oscillations in amplitude and phase domains (Buzsaki, 2015).

To the best of our knowledge there are no studies that have evaluated cognitive processing as a function of respiratory phase. However, interactions between respiration and non-respiratory functions have been documented in humans and rodents. In humans, for example, phase-locking with respiration has been observed for eye movements (Rassler and Raabe, 2003; Rittweger and Popel, 1998), finger movements (Ebert et al., 2002; Rassler, 2000; Rassler et al., 1996) and grip-force (Li and Laskin, 2006). In mice, movements of the mystacial whiskers are phase-locked to respiration (Cao et al., 2012; Moore et al., 2013).

Respiration has also been implicated in the modulation of pain perception. Pain-studies in humans showed that pain perception is reduced during inspiration (Arsenault et al., 2013) and that focused slow breathing reduces the perceived severity of pain (Zautra et al., 2010). Other clinical studies have shown that strength of cortico-spinal communication assessed with trans-cranial magnetic stimulation (TMS) is modulated in phase with respiration (Li and Rymer, 2011). In view of our findings we suggest that these interactions between respiration and sensory-motor processes are, at least in part, caused by respiration-locked fluctuations of ongoing neuronal activity in motor and sensory cortical areas.

In summary, we propose that ongoing neuronal activity of the neocortex is rhythmically modulated by the act of breathing. Respiration-locked neocortical activity is caused by afferent sensory inputs, which convey mechanical and chemical signals associated with every breath. We described three types of cortical activity that are phase-locked to respiration. Two activity types, gamma oscillations and phase transitions have been implicated in cognitive function. Our findings open up new vistas of the relation of cortical neuronal activity to sensory states of the body that are defined by the respiratory cycle. This new physiological role of respiration calls for experimental designs to incorporate respiratory information and for future investigations of the interactions between respiration and cognitive, sensory and motor processes.

## Experimental Procedures

### Animal studies

#### Mice

Experiments were performed on adult male C57BL/6J (B6) mice (> 8 weeks old, 18-25 g body weight). Mice were housed in a breeding colony with 12-hour light/dark cycles in standard cages housing maximally 5 adult mice with ad libitum access to food and water. All experiments were performed during the light cycle (between 12:00 and 17:00 h). None of the mice had undergone any previous experimental procedure. All animal experimental procedures adhered to guidelines approved by the University of Tennessee Health Science Center Animal Care and Use Committee. Principles of laboratory animal care (NIH publication No. 86-23, rev. 1996) were followed.

#### Surgical preparation for awake, head fixed recording

For surgeries mice were initially anesthetized with 3% Isoflurane (Baxter Pharmaceutical Products, Deerfield IL) in oxygen in an incubation chamber, transferred to a stereotaxic head mount and anesthesia was continued with 1-2.5% Isoflurane in oxygen through a mouthpiece. Isoflurane concentration was controlled with a vaporizer (Highland Medical Equipment, CA). The depth of anesthesia was adjusted such that mice failed to show a reflex withdrawal of the hind paw to a strong pinch. Blunt ear bars were used to prevent damaging of the eardrums. Core body temperature, measured with a rectal thermometer, was maintained between 36.5 and 38.0°C with a feedback controlled heating pad (FHC Inc., Bowdoinham, ME). Surgical techniques were described in detail elsewhere (Bryant et al., 2010; Bryant et al., 2009). In brief, a small craniotomy (diameter, 1–2 mm) was made over the target region of cortex or the olfactory bulb. The exposed but intact dura was covered with Triple Antibiotic (Walgreens, US) to keep it moist and reduce the risk of infection. A cylindrical plastic chamber (0.45 cm diameter and 8 mm height) was placed over the skull opening and filled with Triple Antibiotic. Three small machine screws (1/8’ dome head, 0.8 mm diameter, 2 mm long, Small Parts, Inc., Miami Lakes, FL) were secured in the skull bone and metal head post was mounted anterior to Bregma. The chamber, head post and skull screws were secured into place with dental acrylic, as described earlier. Mice were injected subcutaneously with an analgesic (0.05ml Carprofen) to alleviate pain and aid recovery.

#### Electrophysiological recordings and respiration monitoring in awake mice

Mice were allowed a 3–4-day recovery period after surgical mounting of a head post and recording chamber before recordings sessions started. During these sessions the head was held fixed and the body was covered with a loose fitting plastic half-tube (5 cm diameter, 10 cm long) to limit movements. Mice typically adapted to the head fixation within 30 – 45 min as judged by markedly reduced walking and running movements. The experimental setup, head-holding device and recording procedures have been described in detail in a technical publication (Bryant et al., 2009). In short, head fixation involved clamping the metal head-post mounted on the mouse’s skull to a metal fixation device, using a machine screw. Triple antibiotic paste was then removed from the recording chamber and the chamber rinsed and filled with sterile saline solution. Before each experiment, mice were allowed to accommodate to the head fixation situation for 30 min prior to recordings.

For extracellular recordings the guiding tubes of a computer-controlled microdrive (MiniMatrix, Thomas Recording, Germany) were lowered into the saline-filled recording chamber to a distance of less than 2 mm from the dural surface of the brain. The stainless steel guiding tubes also serve as reference electrodes and are eclectically connected to the brain tissue via the saline solution. Then, 2-5 electrodes (glass insulated tungsten/platinum, impedance: 3.5-5.0 MΩ) were slowly advanced through the intact dura into the whisker barrel cortex. Electrode movements were controlled with micrometer resolution and digitally monitored. Local field potentials and spike signals were separated by band pass filtering at 0.1 to 200 Hz and at 200 Hz to 8 kHz, respectively, using a hardware filter amplifier (FA32; Multi Channel Systems). Filtered and amplified voltage signals were digitized and stored on a computer hard disk (16 bit A/D converter; sampling rate, >20 kHz for action potentials, >1 kHz for LFPs) using a CED power1401 and Spike2 software (both Cambridge Electronic Design).

Respiratory behavior was monitored based on temperature changes associated with the expiration of warm air. A thermistor (Measurement Specialties Inc., Boston, MA, USA) was placed in front of one nostril (or descending trachea in some tracheotomy experiments) and breathing cycles could reliably be measured as temperature increased and decreased during exhale and inhale movements respectively. Increased temperature is represented as positive deflections of the thermistor voltage signal, which thus correspond to exhale movements (Figs. 1 and 2). The raw thermistor voltage signal was digitized at 1 kHz and stored together with the electrophysiological signals.

Upon completion of each recording session the Ringer’s solution was removed, the recording chamber filled with triple antibiotic and the mice were returned to their home cages. Each animal typically participated in experiments for 1-2 weeks.

#### Analysis of local field potentials

LFP data were collected in awake, head-fixed conditions from 5 intact and 6 olfactory bulbectomized mice. Primary analysis involved identification of average shape of LFP modulation by respiration by calculating respiration-triggered LFP averages and the strength of phase locking between LFP oscillations and respiration using coherence analysis. Periods of stable resting respiratory rhythm (<5Hz) were selected for further analysis, excluding periods of higher respiratory rates associated with sniffing.

Respiration triggered averages of LFP activity were calculated by aligning LFP signals on inspiration onset times. Whether LFP activity was significantly modulated in phase with respiration was determined using bootstrap statistics (Diaconis and Efron, 1983; Efron and Tibshirani, 1993). To this end, inspiration-onset time markers were randomly shifted in time and a new LFP average was calculated. This shuffling of time-markers and recalculating was repeated >100 times and the resulting population of surrogate LFP averages were rank-ordered and the 95 and 5%ile boundaries of the surrogate distribution were determined. If the original LFP average exceeded either boundary we concluded that the LFP activity was significantly modulated in phase with respiration.

#### Analysis of spiking activity

Single unit spiking activity was recorded together with LFPs. The high frequency spike component (300 – 8000Hz) and low frequency LFP component (0.1 – 200Hz) were separated by band pass filters prior to sampling the two signals in separate channels. Spike events were extracted using a fixed threshold and then sorted based on spike shape using an off-line sorting algorithm (Plexon Inc, Dallas, TX). If the signal to noise ratio of a spike signal was larger than 4 and the inter-spike-intervals of sorted spike trains had a gamma distribution with a refractory period >5 ms we regarded it as a single unit. From the respiratory signals we extracted the times of voltage minima, which correspond to the end of expiration. Cross-correlation analysis of spike times and times of inspiration onsets was performed to determine whether spike activity was modulated with the rhythm of respiration. To determine the significance of spike-respiration correlations we generated surrogate correlations by randomly shifting the spike train in time relative to the respiratory times. The spike train could be shifted in time by any random value between 0 sec and the full duration of the recording. All spike times were shifted by the same value, preserving the spike interval sequence. The process was repeated >100 times, each time calculating the correlation of the shifted spike trains and saving the correlation values. From these >100 surrogate correlations we calculated the median and the 5^th^ and 95^th^ percentile of the distributions of correlation coefficients at each lag time. Correlations were considered significant if the raw correlation coefficient values exceeded the 5th or 95th percentile of the surrogate distributions (Fig. 2).

### Human Studies

Patients had intracranial electrodes placed as part of their individual epilepsy surgery evaluation. In some patients the electrodes were used to precisely define the epileptogenic zone prior to surgical removal, in others also for functional mapping of eloquent cortex prior to surgery. Patients then underwent continuous video-ECoG recordings to capture their typical seizures and to perform functional mapping, if necessary. Recordings were typically done for 5-7 days. ECoG signals were band-pass filtered (0.1 Hz - 70 Hz) prior to digitization at a sampling rate of 1 kHz.

Patients wore a respiratory (stretch-sensitive) band around their chest to determine respiratory status. The respiratory signal was digitized and registered together with ECoG signals. Inspiration (belt stretching) resulted in an increase of a voltage signal. The time of transition from voltage increase to decrease (time of peak voltage) thus corresponded to the time of transition from inspiration to expiration. This uniquely defined feature of the respiration-voltage signal was used as a temporal align for further analysis of respiration-locked features of ECoG activity. The ECoG/respiratory data were collected from quiet periods on the third day or later after placement of intracranial electrodes.

## Acknowledgements

This research was supported by a grant from the UTHSC College of Medicine iRISE Pilot Program to DHH, ACP and JWW and by support from the Dept. Anatomy & Neurobiology, Univ. Tennessee Health Sci. Center to DHH. The contribution by RK has been supported in part by NSF CRCNS grant NSF-DMS-13-11165. SSM was supported by the UTHSC Neuroscience Institute.

## References

Adrian, E.D. (1950). The electrical activity of the mammalian olfactory bulb. Electroencephalography and clinical neurophysiology 2, 377–388.

Arieli, A., Sterkin, A., Grinvald, A., and Aertsen, A. (1996). Dynamics of ongoing activity: Explanation of the large variability in evoked cortical responses. Science 273, 1868–1871.

Arsenault, M., Ladouceur, A., Lehmann, A., Rainville, P., and Piche, M. (2013). Pain modulation induced by respiration: phase and frequency effects. Neuroscience 252, 501–511.

Barbour, A.D., Holst, L., and Janson, S. (1992). Poisson Approximation (Oxford: Clarendon Press).

Belvisi, M.G. (2003). Airway sensory innervation as a target for novel therapies: an outdated concept? Current opinion in pharmacology 3, 239–243.

Boly, M., Balteau, E., Schnakers, C., Degueldre, C., Moonen, G., Luxen, A., Phillips, C., Peigneux, P., Maquet, P., and Laureys, S. (2007). Baseline brain activity fluctuations predict somatosensory perception in humans. Proc Natl Acad Sci U S A 104, 12187–12192.

Bonifazi, P., Goldin, M., Picardo, M.A., Jorquera, I., Cattani, A., Bianconi, G., Represa, A., Ben-Ari, Y., and Cossart, R. (2009). GABAergic Hub Neurons Orchestrate Synchrony in Developing Hippocampal Networks. Science 326, 1419–1424.

Bryant, J.L., Boughter, J.D., Gong, S., LeDoux, M.S., and Heck, D.H. (2010). Cerebellar cortical output encodes temporal aspects of rhythmic licking movements and is necessary for normal licking frequency. Eur J Neurosci 32, 41–52.

Bryant, J.L., Roy, S., and Heck, D.H. (2009). A technique for stereotaxic recordings of neuronal activity in awake, head-restrained mice. Journal of Neuroscience Methods 178, 75.

Buzsaki, G. (2015). Hippocampal sharp wave-ripple: A cognitive biomarker for episodic memory and planning. Hippocampus 25, 1073–1188.

Canolty, R.T., Edwards, E., Dalal, S.S., Soltani, M., Nagarajan, S.S., Kirsch, H.E., Berger, M.S., Barbaro, N.M., and Knight, R.T. (2006). High gamma power is phase-locked to theta oscillations in human neocortex. Science 313, 1626–1628.

Cao, Y., Roy, S., Sachdev, R.N., and Heck, D.H. (2012). Dynamic correlation between whisking and breathing rhythms in mice. J Neurosci 32, 1653–1659.

Cape, E.G., and Jones, B.E. (2000). Effects of glutamate agonist versus procaine microinjections into the basal forebrain cholinergic cell area upon gamma and theta EEG activity and sleep-wake state. Eur J Neurosci 12, 2166–2184.

Carstens, E., Leah, J., Lechner, J., and Zimmermann, M. (1990). Demonstration of extensive brainstem projections to medial and lateral thalamus and hypothalamus in the rat. Neuroscience 35, 609–626.

Chauvette, S., Seigneur, J., and Timofeev, I. (2012). Sleep oscillations in the thalamocortical system induce long-term neuronal plasticity. Neuron 75, 1105–1113.

Chen, Z., Eldridge, F.L., and Wagner, P.G. (1992). Respiratory-associated thalamic activity is related to level of respiratory drive. Respiration physiology 90, 99–113.

Craig, A.D. (2002). How do you feel? Interoception: the sense of the physiological condition of the body. Nat Rev Neurosci 3, 655–666.

Craig, A.D. (2009). How do you feel–now? The anterior insula and human awareness. Nat Rev Neurosci 10, 59–70.

Csicsvari, J., Hirase, H., Czurko, A., Mamiya, A., and Buzsaki, G. (1999). Fast network oscillations in the hippocampal CA1 region of the behaving rat. JNeurosci 19, RC20.

Davenport, P.W., and Vovk, A. (2009). Cortical and subcortical central neural pathways in respiratory sensations. Respiratory physiology & neurobiology 167, 72–86.

Deco, G., Jirsa, V.K., and McIntosh, A.R. (2011). Emerging concepts for the dynamical organization of resting-state activity in the brain. Nat Rev Neurosci 12, 43–56.

Diaconis, P., and Efron, B. (1983). Computer-Intensive Methods in Statistics. SciAm 248, 116–130.

Dillbeck, M.C., and Bronson, E.C. (1981). Short-term longitudinal effects of the transcendental meditation technique on EEG power and coherence. Int J Neurosci 14, 147–151.

Ebert, D., Hefter, H., Binkofski, F., and Freund, H.J. (2002). Coordination between breathing and mental grouping of pianistic finger movements. Percept Mot Skills 95, 339–353.

Eeckman, F.H., and Freeman, W.J. (1990). Correlations between unit firing and EEG in the rat olfactory system. Brain Res 528, 238–244.

Efron, B., and Tibshirani, R.J. (1993). An Introduction to the Bootstrap, Vol 57 (Washington, D.C.: Chapman & Hall/CRC).

Engel, A.K., Fries, P., and Singer, W. (2001). Dynamic predictions: oscillations and synchrony in top-down processing. NatRevNeurosci 2, 704–716.

Erdős, P., and Rényi, A. (1960). On the evolution of random graphs. Magyar Tudoamányos Akadémia, Mat Kut Int Közl 5, 17–61.

Feng, W., Stormer, V.S., Martinez, A., McDonald, J.J., and Hillyard, S.A. (2014). Sounds activate visual cortex and improve visual discrimination. J Neurosci 34, 9817–9824.

Ferezou, I., Haiss, F., Gentet, L.J., Aronoff, R., Weber, B., and Petersen, C.C. (2007). Spatiotemporal dynamics of cortical sensorimotor integration in behaving mice. Neuron 56, 907–923.

Finger, T.E., Bottger, B., Hansen, A., Anderson, K.T., Alimohammadi, H., and Silver, W.L. (2003). Solitary chemoreceptor cells in the nasal cavity serve as sentinels of respiration. Proc Natl Acad Sci U S A 100, 8981–8986.

Fontanini, A., and Bower, J.M. (2005). Variable coupling between olfactory system activity and respiration in ketamine/xylazine anesthetized rats. Journal of Neurophysiology 93, 3573–3581.

Fontanini, A., and Bower, J.M. (2006). Slow-waves in the olfactory system: an olfactory perspective on cortical rhythms. Trends in neurosciences 29, 429–437.

Fontanini, A., Spano, P., and Bower, J.M. (2003). Ketamine-xylazine-induced slow (< 1.5 Hz) oscillations in the rat piriform (olfactory) cortex are functionally correlated with respiration. Journal of Neuroscience 23, 7993–8001.

Fox, M.D., and Raichle, M.E. (2007). Spontaneous fluctuations in brain activity observed with functional magnetic resonance imaging. Nat Rev Neurosci 8, 700–711.

Freeman, W.J. (1975). Mass action in the nervous system (New York: Academic Press).

Freeman, W.J. (2004a). Origin, structure, and role of background EEG activity. Part 1. Analytic amplitude. Clinical neurophysiology: official journal of the International Federation of Clinical Neurophysiology 115, 2077–2088.

Freeman, W.J. (2004b). Origin, structure, and role of background EEG activity. Part 2. Analytic phase. Clinical neurophysiology: official journal of the International Federation of Clinical Neurophysiology 115, 2089–2107.

Freeman, W.J. (2009). Deep analysis of perception through dynamic structures that emerge in cortical activity from self-regulated noise. Cogn Neurodyn 3, 105–116.

Freeman, W.J. (2015). Mechanism and significance of global coherence in scalp EEG. Curr Opin Neurobiol 31, 199–205.

Freeman, W.J., Holmes, M.D., West, G.A., and Vanhatalo, S. (2006). Fine spatiotemporal structure of phase in human intracranial EEG. Clinical neurophysiology: official journal of the International Federation of Clinical Neurophysiology 117, 1228–1243.

Freeman, W.J., Kozma, R., Bollobas, B., and Riordan, O. (2009). Scale-free cortical planar networks. In Handbook of Large-Scale Random Networks (Berlin: Springer), pp. 277–324.

Freeman, W.J., and Rogers, L.J. (2002). Fine temporal resolution of analytic phase reveals episodic synchronization by state transitions in gamma EEGs. J Neurophysiol 87, 937–945.

Fries, P., Reynolds, J.H., Rorie, A.E., and Desimone, R. (2001). Modulation of oscillatory neuronal synchronization by selective visual attention. Science 291, 1560–1563.

Gallos, L.K., Makse, H.A., and Sigman, M. (2012). A small world of weak ties provides optimal global integration of self-similar modules in functional brain networks. Proceedings of the National Academy of Sciences of the United States of America 109, 2825–2830.

Gaylord, C., Orme-Johnson, D., and Travis, F. (1989). The effects of the transcendental mediation technique and progressive muscle relaxation on EEG coherence, stress reactivity, and mental health in black adults. Int J Neurosci 46, 77–86.

Georgopoulos, A.P., Schwartz, A.B., and Kettner, R.E. (1986). Neuronal Population Coding of Movement Direction. Science 233, 1416–1419.

Gilad, Y., Przeworski, M., and Lancet, D. (2004). Loss of olfactory receptor genes coincides with the acquisition of full trichromatic vision in primates. PLoS Biol 2, E5.

Gould, I.C., Nobre, A.C., Wyart, V., and Rushworth, M.F. (2012). Effects of decision variables and intraparietal stimulation on sensorimotor oscillatory activity in the human brain. J Neurosci 32, 13805–13818.

Grosmaitre, X., Santarelli, L.C., Tan, J., Luo, M., and Ma, M. (2007). Dual functions of mammalian olfactory sensory neurons as odor detectors and mechanical sensors. Nature neuroscience 10, 348–354.

He, B.J. (2013). Spontaneous and task-evoked brain activity negatively interact. J Neurosci 33, 4672–4682.

Herrmann, C.S., Frund, I., and Lenz, D. (2010). Human gamma-band activity: a review on cognitive and behavioral correlates and network models. Neurosci Biobehav Rev 34, 981–992.

Honey, C.J., Thivierge, J.P., and Sporns, O. (2010). Can structure predict function in the human brain? Neuroimage 52, 766–776.

Immanuel, S.A., Pamula, Y., Kohler, M., Martin, J., Kennedy, D., Saint, D.A., and Baumert, M. (2014). Respiratory cycle-related electroencephalographic changes during sleep in healthy children and in children with sleep disordered breathing. Sleep 37, 1353–1361.

Ito, J., Maldonado, P., and Grun, S. (2013). Cross-frequency interaction of the eye-movement related LFP signals in V1 of freely viewing monkeys. Front Syst Neurosci 7, 1.

Ito, J., Roy, S., Liu, Y., Cao, Y., Fletcher, M., Boughter, J.D., Jr., Grun, S., and Heck, D.H. (2014). Whisker barrel cortex delta oscillations and gamma power in the awake mouse are linked to respiration. Nat Commun 5.

Janson, S., Kozma, R., Ruszinkó, M., and Sokolov, Y. (2016). Bootstrap percolation on a random graph coupled with a lattice. submitted.

Kay, L.M. (2015). Olfactory system oscillations across phyla. Curr Opin Neurobiol 31, 141–147.

Kay, L.M., and Beshel, J. (2010). A beta oscillation network in the rat olfactory system during a 2-alternative choice odor discrimination task. J Neurophysiol 104, 829–839.

Kay, L.M., Beshel, J., Brea, J., Martin, C., Rojas-Libano, D., and Kopell, N. (2009). Olfactory oscillations: the what, how and what for. Trends Neurosci 32, 207–214.

Kay, L.M., and Freeman, W.J. (1998). Bidirectional processing in the olfactory-limbic axis during olfactory behavior. Behav Neurosci 112, 541–553.

Kim, D.J., Bolbecker, A.R., Howell, J., Rass, O., Sporns, O., Hetrick, W.P., Breier, A., and ODonnell, B.F. (2013). Disturbed resting state EEG synchronization in bipolar disorder: A graph-theoretic analysis. Neuroimage-Clin 2, 414–423.

Kozma, R., Davis, J.J., and Freeman, W.J. (2012). Synchronization of de-synchronization events demonstrate large-scale cortical singularities as hallmarks of higher cognitive activity. J Neurosci Neuro-Engng 1, 13–23.

Kozma, R., and Freeman, W.J. (2001). Analysis of visual theta rhythm-experimental and theoretical evidence of visual sniffing. IJCNN01 2, 1118–1121.

Kozma, R., and Freeman, W.J. (2008). Intermittent spatio-temporal desynchronization and sequenced synchrony in ECoG signals. Chaos 18, 037131.

Kozma, R., and Freeman, W.J. (2009). The KIV model of intentional dynamics and decision making. Neural Networks 22, 277–285.

Kozma, R., and Freeman, W.J. (2016). Cognitive Phase Transitions in the Cerebral Cortex-Enhancing the Neuron Doctrine by Modeling Neural Fields (Springer International Publishing).

Kozma, R., and Puljic, M. (2015). Random graph theory and neuropercolation for modeling brain oscillations at criticality. Curr Opin Neurobiol 31, 181–188.

Kozma, R., Ruszinkó, M., and Sokolov, Y. (2015). Activation process on a long-range percolation graph with power law long edge distribution. Part 2: Two types of vertices. (In Preparation).

Kozma, R., Ruszinkó, M., and Sokolov, Y. (2016). Bootstrap percolation on a random graph coupled with a lattice. Part II: Two types of vertices. in progress.

Krout, K.E., Belzer, R.E., and Loewy, A.D. (2002). Brainstem projections to midline and intralaminar thalamic nuclei of the rat. J Comp Neurol 448, 53–101.

Laufs, H., Krakow, K., Sterzer, P., Eger, E., Beyerle, A., Salek-Haddadi, A., and Kleinschmidt, A. (2003). Electroencephalographic signatures of attentional and cognitive default modes in spontaneous brain activity fluctuations at rest. Proc Natl Acad Sci U S A 100, 11053–11058.

Lee, C., Rohrer, W.H., and Sparks, D.L. (1988). Population coding of saccadic eye movements by neurons in the superior colliculus. Nature 332, 357–360.

Li, S., Bai, W., Liu, T., Yi, H., and Tian, X. (2012). Increases of theta-low gamma coupling in rat medial prefrontal cortex during working memory task. Brain research bulletin 89, 115–123.

Li, S., and Laskin, J.J. (2006). Influences of ventilation on maximal isometric force of the finger flexors. Muscle & nerve 34, 651–655.

Li, S., and Rymer, W.Z. (2011). Voluntary breathing influences corticospinal excitability of nonrespiratory finger muscles. J Neurophysiol 105, 512–521.

Linkenkaer-Hansen, K., Nikulin, V.V., Palva, S., Ilmoniemi, R.J., and Palva, J.M. (2004). Prestimulus oscillations enhance psychophysical performance in humans. J Neurosci 24, 10186–10190.

Linster, C., and Hasselmo, M.E. (2000). Neural activity in the horizontal limb of the diagonal band of broca can be modulated by electrical stimulation of the olfactory bulb and cortex in rats. Neurosci Lett 282, 157–160.

Liotti, M., Brannan, S., Egan, G., Shade, R., Madden, L., Abplanalp, B., Robillard, R., Lancaster, J., Zamarripa, F.E., Fox, P.T., and Denton, D. (2001). Brain responses associated with consciousness of breathlessness (air hunger). Proc Natl Acad Sci USA 98, 2035–2040.

Lorig, T.S., Schwartz, G.E., and Herman, K.B. (1988). Brain and odor: II. EEG activity during nose and mouth breathing. Psychobiology 16, 285–287.

Lowet, E., Roberts, M.J., Bosman, C.A., Fries, P., and de Weerd, P. (2015). Areas V1 and V2 show microsaccade-related 3-4 Hz covariation in gamma power and frequency. Eur J Neurosci.

Macey, P.M., Woo, M.A., Macey, K.E., Keens, T.G., Saeed, M.M., Alger, J.R., and Harper, R.M. (2005). Hypoxia reveals posterior thalamic, cerebellar, midbrain, and limbic deficits in congenital central hypoventilation syndrome. JApplPhysiol 98, 958–969.

Manns, I.D., Alonso, A., and Jones, B.E. (2003). Rhythmically discharging basal forebrain units comprise cholinergic, GABAergic, and putative glutamatergic cells. J Neurophysiol 89, 1057–1066.

Marshall, L., Helgadottir, H., Molle, M., and Born, J. (2006). Boosting slow oscillations during sleep potentiates memory. Nature 444, 610–613.

Martinez-Conde, S., Otero-Millan, J., and Macknik, S.L. (2013). The impact of microsaccades on vision: towards a unified theory of saccadic function. Nat Rev Neurosci 14, 83–96.

Mason, P., Gao, K., and Genzen, J.R. (2007). Serotonergic raphe magnus cell discharge reflects ongoing autonomic and respiratory activities. J Neurophysiol 98, 1919–1927.

Maynard, E.M., Hatsopoulos, N.G., Ojakangas, C.L., Acuna, B.D., Sanes, J.N., Normann, R.A., and Donoghue, J.P. (1999). Neuronal Interactions Improve Cortical Population Coding of Movement Direction. Journal of Neuroscience 19, 8083–8093.

McDonald, J.J., Stormer, V.S., Martinez, A., Feng, W., and Hillyard, S.A. (2013). Salient sounds activate human visual cortex automatically. J Neurosci 33, 9194–9201.

Moore, J.D., Deschenes, M., Furuta, T., Huber, D., Smear, M.C., Demers, M., and Kleinfeld, D. (2013). Hierarchy of orofacial rhythms revealed through whisking and breathing. Nature 497, 205–210.

Nacher, V., Ledberg, A., Deco, G., and Romo, R. (2013). Coherent delta-band oscillations between cortical areas correlate with decision making. Proc Natl Acad Sci U S A 110, 15085–15090.

Nguyen Chi, V., Muller, C., Wolfenstetter, T., Yanovsky, Y., Draguhn, A., Tort, A.B., and Brankack, J. (2016). Hippocampal Respiration-Driven Rhythm Distinct from Theta Oscillations in Awake Mice. J Neurosci 36, 162–177.

Niell, C.M., and Stryker, M.P. (2010). Modulation of visual responses by behavioral state in mouse visual cortex. Neuron 65, 472–479.

Palva, J.M., Zhigalov, A., Hirvonen, J., Korhonen, O., Linkenkaer-Hansen, K., and Palva, S. (2013). Neuronal long-range temporal correlations and avalanche dynamics are correlated with behavioral scaling laws. Proc Natl Acad Sci U S A 110, 3585–3590.

Phillips, M.E., Sachdev, R.N., Willhite, D.C., and Shepherd, G.M. (2012). Respiration drives network activity and modulates synaptic and circuit processing of lateral inhibition in the olfactory bulb. Journal of Neuroscience 32, 85–98.

Poulet, J.F., and Petersen, C.C. (2008). Internal brain state regulates membrane potential synchrony in barrel cortex of behaving mice. Nature 454, 881–885.

Rassler, B. (2000). Mutual nervous influences between breathing and precision finger movements. European journal of applied physiology 81, 479–485.

Rassler, B., Ebert, D., Waurick, S., and Junghans, R. (1996). Coordination Between Breathing and Finger Tracking in Man. Journal of motor behavior 28, 48–56.

Rassler, B., and Raabe, J. (2003). Co-ordination of breathing with rhythmic head and eye movements and with passive turnings of the body. European journal of applied physiology 90, 125–130.

Reijneveld, J.C., Ponten, S.C., Berendse, H.W., and Stam, C.J. (2007). The application of graph theoretical analysis to complex networks in the brain. Clinical Neurophysiology 118, 2317–2331.

Ritter, S.M., Strick, M., Bos, M.W., van Baaren, R.B., and Dijksterhuis, A. (2012). Good morning creativity: task reactivation during sleep enhances beneficial effect of sleep on creative performance. Journal of sleep research 21, 643–647.

Rittweger, J., and Popel, A. (1998). Respiratory-like periodicities in slow eye movements during sleep onset. Clinical physiology 18, 471–478.

Rowe, T.B., Macrini, T.E., and Luo, Z.X. (2011). Fossil evidence on origin of the mammalian brain. Science 332, 955–957.

Sadaghiani, S., Hesselmann, G., and Kleinschmidt, A. (2009). Distributed and antagonistic contributions of ongoing activity fluctuations to auditory stimulus detection. J Neurosci 29, 13410–13417.

Schelegle, E.S. (2003). Functional morphology and physiology of slowly adapting pulmonary stretch receptors. The anatomical record Part A, Discoveries in molecular, cellular, and evolutionary biology 270, 11–16.

Schölvinck, M.L., Saleem, A.B., Benucci, A., Harris, K.D., and Carandini, M. (2015). Cortical state determines global variability and correlations in visual cortex. J Neurosci 35, 170–178.

Servit, Z., Kristof, M., and Kolinova, M. (1977). Activation of epileptic electrographic phenomena in the human EEG by nasal air flow. Physiol Bohemoslov 26, 499–506.

Shadlen, M.N., and Newsome, W.T. (1994). Noise, neural codes and cortical organization. Current Opinion in Neurobiology 4, 569–579.

Sheth, B.R., Sandkuhler, S., and Bhattacharya, J. (2009). Posterior Beta and anterior gamma oscillations predict cognitive insight. J Cogn Neurosci 21, 1269–1279.

Siegel, M., Engel, A.K., and Donner, T.H. (2011). Cortical network dynamics of perceptual decision-making in the human brain. Front Hum Neurosci 5, 21.

Sporns, O., Tononi, G., and Kotter, R. (2005). The human connectome: A structural description of the human brain. Plos Computational Biology 1, 245–251.

Stam, C.J., Jones, B.F., Nolte, G., Breakspear, M., and Scheltens, P. (2007). Small-world networks and functional connectivity in Alzheimer’s disease. Cerebral Cortex 17, 92–99.

Stancak, A., Jr., and Kuna, M. (1994). EEG changes during forced alternate nostril breathing. Int J Psychophysiol 18, 75–79.

Tallon-Baudry, C. (2003). Oscillatory synchrony and human visual cognition. J Physiol Paris 97, 355–363.

Tallon-Baudry, C. (2004). Attention and awareness in synchrony. Trends Cogn Sci 8, 523–525.

Tort, A.B., Komorowski, R.W., Manns, J.R., Kopell, N.J., and Eichenbaum, H. (2009). Theta-gamma coupling increases during the learning of item-context associations. Proc Natl Acad Sci 106, 20942–20947.

Tsanov, M., Chah, E., Reilly, R., and OMara, S.M. (2014). Respiratory cycle entrainment of septal neurons mediates the fast coupling of sniffing rate and hippocampal theta rhythm. Eur J Neurosci 39, 957–974.

Turova, T.S., and Villa, A.E.P. (2007). On a phase diagram for random neural networks with embedded spike timing dependent plasticity. Biosystems 89, 280–286.

van Vugt, M.K., Simen, P., Nystrom, L.E., Holmes, P., and Cohen, J.D. (2012). EEG oscillations reveal neural correlates of evidence accumulation. Front Neurosci 6, 106.

Vinnik, E., Itskov, P.M., and Balaban, E. (2012). beta-And gamma-band EEG power predicts illusory auditory continuity perception. J Neurophysiol 108, 2717–2724.

Widdicombe, J. (2009). Lung afferent activity: implications for respiratory sensation. Respiratory physiology & neurobiology 167, 2–8.

Wyart, V., de Gardelle, V., Scholl, J., and Summerfield, C. (2012). Rhythmic fluctuations in evidence accumulation during decision making in the human brain. Neuron 76, 847–858.

Yanovsky, Y., Ciatipis, M., Draguhn, A., Tort, A.B., and Brankack, J. (2014). Slow oscillations in the mouse hippocampus entrained by nasal respiration. J Neurosci 34, 5949–5964.

Zautra, A.J., Fasman, R., Davis, M.C., and Craig, A.D. (2010). The effects of slow breathing on affective responses to pain stimuli: an experimental study. Pain 149, 12–18.

